# Energy Shortage in Human and Mouse Models of *SLC4A11*-Associated Corneal Endothelial Dystrophies

**DOI:** 10.1101/868281

**Authors:** Wenlin Zhang, Ricardo Frausto, Doug D. Chung, Christopher G. Griffis, Liyo Kao, Angela Chen, Rustam Azimov, Alapakkam P. Sampath, Ira Kurtz, Anthony J. Aldave

## Abstract

**Purpose:** To elucidate the molecular events in solute carrier family 4 member 11 (SLC4A11)-deficient corneal endothelium that lead to the endothelial dysfunction that characterizes the dystrophies associated with *SLC4A11* mutations, congenital hereditary endothelial dystrophy (CHED) and Fuchs endothelial corneal dystrophy 4.

**Methods:** Comparative transcriptomic analysis (CTA) was performed in primary human corneal endothelial cells (pHCEnC) and murine corneal endothelial cells (MCEnC) with normal and reduced levels of SLC4A11 (*SLC4A11* KD pHCEnC) and Slc4a11 (*Slc4a11^−/−^* MCEnC), respectively. Validation of differentially expressed genes was performed using immunofluorescence staining of CHED corneal endothelium, as well as western blot and quantitative PCR analysis of *SLC4A11* KD pHCEnC and *Slc4a11^−/−^* MCEnC. Functional analyses were performed to investigate potential functional changes associated with the observed transcriptomic alterations.

**Results:** CTA revealed inhibition of cell metabolism and ion transport function as well as mitochondrial dysfunction, leading to reduced adenosine triphosphate (ATP) production, in *SLC4A11* KD pHCEnC and *Slc4a11^−/−^* MCEnC. Co-localization of SNARE protein STX17 with mitochondria marker COX4 was observed in CHED corneal endothelium, as was activation of AMPK–p53/ULK1 in both *SLC4A11* KD pHCEnC and *Slc4a11^−/−^* MCEnC, providing additional evidence of mitochondrial dysfunction and mitophagy. Reduced Na^+^-dependent HCO_3_^−^ transport activity and altered NH_4_Cl-induced membrane potential changes were observed in *Slc4a11^−/−^* MCEnC.

**Conclusions:** Reduced steady-state ATP levels and subsequent activation of the AMPK–p53 pathway provide a link between the metabolic functional deficit and transcriptome alterations, as well as evidence of insufficient ATP to maintain the Na^+^/K^+^-ATPase corneal endothelial pump as the cause of the edema that characterizes *SLC4A11*-associated corneal endothelial dystrophies.

---

Solute carrier family 4 member 11 (SLC4A11) is one of the highly expressed differentiation markers for corneal endothelium.^1–3^ Mutations in *SLC4A11* are associated with congenital hereditary endothelial dystrophy (CHED), Harboyan syndrome (CHED with perceptive deafness) and a subset of Fuchs endothelial corneal dystrophy (FECD4).^4–7^ Children with CHED often present with bilateral corneal edema at or shortly after birth with significant vision impairment. Corneal transplantation is the only means of restoring vision and is associated with a guarded prognosis in terms of graft survival and long-term recovery of vision.^8^ In addition, these children are at risk of developing perceptive deafness later in life (Harboyan syndrome).^9^ FECD affects as many as 5% of the US population over 40 years of age,^10^ and visually significant corneal edema secondary to FECD is the most common indication for keratoplasty in the United States and worldwide.^11,12^ Together, CHED and FECD constitute common indications for corneal transplantation in published series from around the world.^13,14^

SLC4A11 is functionally characterized as an NH3 and alkaline pH-stimulated H^+^ transporter, while permeability to Na^+^, OH^−^ and water has also been reported.^15–19^ SLC4A11 is essential in facilitating energy-producing glutaminolysis, maintaining antioxidant signaling, and preventing apoptosis in corneal endothelial cells (CEnC).^20–23^ During development and in the event of oxidative DNA damage, *SLC4A11* gene expression is upregulated by direct binding of phosphorylated (activated) p53.^24^ In SLC4A11-associated corneal endothelial dystrophies, the corneal edema that develops as a result of pathologic *SLC4A11* mutations is evidence of CEnC dysfunction, either from direct cell loss/death or from disturbances in the CEnC “pump–leak” system.^25–27^ Corneal transparency is maintained by the CEnC “pump–leak” system through a dynamic balance between the passive leak of aqueous humor fluid from the anterior chamber into the corneal stroma and the active pumping of fluid from the corneal stroma into the anterior chamber. The fluid pump activity is driven by an ionic electrochemical gradient set up by the highly expressed Na^+^/K^+^-ATPase.^28^ As such, CEnC have the second highest density of mitochondria among any cell types in the body (second to photoreceptors) to generate sufficient adenosine triphosphate (ATP) to fuel the Na^+^/K^+^- ATPase-driven endothelial pump.^29^ As SLC4A11 plays a significant role in facilitating ATP-generating glutaminolysis in CEnC,^22^ it is not surprising that glutaminolysis inhibition, mitochondria membrane potential depolarization, enriched mitochondrial reactive oxidative species (ROS), and increased mitochondria turnover have been observed in the CEnC of the *Slc4a11^−/−^* mouse.^22,23,30^ Thus, the association between SLC4A11 and CEnC mitochondrial function suggests that SLC4A11 is involved not only in moving ions across the plasma membrane but also in the supply of energy to the endothelial pump.

Approximately 94 *SLC4A11* mutations have been identified in individuals with CHED.^4,6,9,31–48^ Although a large number of these mutations result in SLC4A11 protein misfolding and failure to mature to the plasma membrane,^5,6,49–51^ some mutations affect SLC4A11 transporter function without impacting membrane trafficking^17,52,53^ or cause aberrant *SLC4A11* pre-mRNA splicing and subsequent reduced SLC4A11 expression.^47^ Collectively, these observations support the hypothesis that loss of SLC4A11 function is the primary pathogenetic mechanism in CHED rather than mutant SLC4A11 protein misfolding/mislocalization in the endoplasmic reticulum (ER). Therefore, we investigated the impact of reduced SLC4A11 function on the CEnC transcriptome in primary human and murine immortalized CEnC, with validation in corneal endothelium from individuals with CHED, and elucidated the upstream molecular mechanism leading to the observed transcriptomic changes.

## Materials and Methods

### Primary Human Corneal Endothelial Cell Culture and Knockdown of SLC4A11

Primary cultures of human corneal endothelial cells (pHCEnC) were established from donor corneas as previously described.^54^ After achieving a confluent monolayer, passage 1 of pHCEnC were transfected with 10 nM anti-SLC4A11 siRNA (CCGAAAGUACCUGAAGUUAAAGAACT) or scrambled siRNA (OriGene Technologies, Rockville, MD, USA) using Lipofectamine LTX (Life Technologies, Carlsbad, CA, USA). At 72 hours post-transfection, the cells were lysed for RNA, protein and ATP isolation.

### Immortalized Mouse Corneal Endothelial Cell Culture

Immortalized *Slc4a11^+/+^* and *Slc4a11^−/−^* mouse corneal endothelial cell (MCEnC) lines were derived from *Slc4a11^+/+^* and *Slc4a11*^−/−^ mice corneal endothelium and were cultured at 33°C in Opti-MEM I medium (Thermo Fisher Scientific, Waltham, MA, USA) with supplements as previously described.^23^ Passages 6 and 39 of *Slc4a11^−/−^* and passages 7 and 40 of *Slc4a11^+/+^* MCEnC were lysed for RNA isolation. Passages 10 and 44 of *Slc4a11^−/−^* and passages 11 and 45 of *Slc4a11^+^*^/*+*^ MCEnC were lysed for protein and ATP isolation.

### Total RNA Isolation from pHCEnC and MCEnC

Total RNA from cultured pHCEnC was isolated in TRI Reagent and purified with the RNeasy Clean-Up Kit (Qiagen, Hilden, Germany). Total RNA from cultured MCEnC was isolated and purified using the Qiagen RNeasy Plus Mini Kit.

### RNA Sequencing of Total RNA from pHCEnC and MCEnC

Purified total RNA from pHCEnC was prepared for RNA sequencing libraries using the KAPA mRNA HyperPrep Kit (Roche Sequencing Solutions, Pleasanton, CA, USA). Libraries were sequenced on the HiSeq 4000 (Illumina, San Diego, CA, USA), and paired-end 150-bp reads were generated. Purified total RNA from the MCEnC was submitted to the UCLA Technology Center for Genomics & Bioinformatics for library preparation and sequencing. Single-end 50-bp reads were generated using the Illumina HiSeq 3000. The generated FASTQ files and quantitative results are available from the NCBI Gene Expression Omnibus database (accession numbers GSE142635 and GSE142636).

### RNA Sequencing Data Analyses

Raw reads from pHCEnC and MCEnC samples were aligned to the human (GRCh38/hg38) and mouse (GRCm38/mm10) genomes, respectively, using HISAT2. Raw counts of aligned reads were converted to counts per million (CPM) mapped reads and normalized by the method of trimmed mean of M-values to adjust for library size differences. Linear models for microarray analysis coupled with variance modeling at the observation level were used for differential gene expression analysis. The CPM fold changes of gene transcripts were calculated by comparing each sample set to the appropriate control: (1) pHCEnC sample set, pHCEnC transfected with siRNA targeting *SLC4A11* (*SLC4A11* KD pHCEnC, *n =* 3) versus pHCEnC transfected with scrambled RNA (scRNA pHCEnC, *n* = 3); (2) MCEnC early sample set, *Slc4a11^−/−^* MCEnC passage 6 (*n* = 4) versus *Slc4a11^+/+^* MCEnC passage 7 (*n =* 4); and (3) MCEnC late sample set, *Slc4a11^−/−^* MCEnC passage 39 (*n* = 4) versus *Slc4a11^+/+^* MCEnC passage 40 (*n* = 4). The following thresholds were applied to define genes with differential expression: CPM > 1, fold change > 1, and adjusted *P* <0.05. A differential gene expression (DGE) list was created for each of the three sample sets, after which comparative transcriptome analysis was performed by comparing the three DGE lists to identify common differentially expressed genes (DEG) and enriched pathways.

### Ingenuity Pathway Analysis

Qiagen Ingenuity Pathway Analysis (IPA) was used to perform comparative transcriptome analysis among the three DGE lists (pHCEnC, MCEnC early, and MCEnC late), including canonical/biological function pathway enrichment and upstream regulator prediction analyses. Enriched canonical and biological function pathways with a predicted activation *z*-score in each of the three DGE lists were sorted by the sign and value of the *z*-score to identify the most enriched pathways. Enriched pathways without an assigned *z*-score and predicted upstream regulators were ranked by mean of enrichment *P* values.

### Quantitative PCR

Quantitative PCR (qPCR) was performed on separate batches of RNA samples isolated from passage 1 of *SLC4A11* KD and scRNA pHCEnC, as well as *Slc4a11^−/−^* MCEnC passages 6 and 39 and of *Slc4a11^+/+^* MCEnC passages 7 and 40. Total RNA of each sample was reversed transcribed using the SuperScript III First-Strand Synthesis System (Sigma-Aldrich, St. Louis, MO, USA). Quantitative PCR was performed on the LightCycler 480 System (Roche, Basel, Switzerland) using Kapa SYBR Fast Universal Kit (Roche) with an annealing temperature of 60°C and with primers listed in Supplementary Table S1. Relative gene expression was calculated by the comparative C_t_ (2^-ΔCt^) method in comparison to the housekeeping gene *PPIA/Ppia* or *ACTB/Actb.*

### Immunofluorescence

Five-micrometer sections of paraffin-embedded corneas from seven healthy donors and two individuals with CHED were de-paraffinized and rehydrated in a graded ethanol series (100%, 95%, 70% and 50%) for 5 minutes each and subject to antigen retrieval in 10 mM sodium citrate. Sections were incubated with primary antibodies overnight at 4°C and with secondary antibodies for 1 hour (Supplementary Table S2). Sections were mounted with Invitrogen ProLong Antifade Mountant (Thermo Fisher Scientific) and imaged with an Olympus FV-1000 inverted confocal fluorescence microscope (Olympus Corporation, Tokyo, Japan). Florescence intensity of the images were quantified using Olympus FluoView 4.2 and ImageJ software (National Institutes of Health, Bethesda, MD, USA).

### Single Cell Patch-Clamp Recordings

Single cell recordings were made from a single adhered MCEnC after culturing of MCEnC on 25-mm glass coverslips to ~10% to 20% confluence. Membrane voltages were measured using whole-cell patch electrodes in current clamp mode (Ihold = 0) as previously described.^55^ Data were reported as mean (95% CI). Details of the experimental setup and solutions used are provided in the Supplementary Materials.

### Intracellular pH Measurement

Intracellular pH (pH_i_) measurements were performed by monitoring intracellular free H^+^ concentration using the pH-sensitive fluorescent dye BCECF-AM (B1170; Thermo Fisher Scientific) as described previously.^19,20^ Details of the experimental setup and solutions used are provided in the Supplementary Materials.

### Western Blotting

Whole-cell lysates from pHCEnC and MCEnC were prepared with radioimmunoprecipitation assay buffer with proteinase and phosphatase inhibitors. Total protein was quantified by bicinchoninic acid (BCA) assay, separated and detected using the Simple Western assay Wes (ProteinSimple, San Jose, CA, USA). Quantification and data analysis were performed in Compass for Simple Western software (Protein-Simple). Antibodies used are listed in Supplementary Table S2.

### Intracellular ATP Assay

*Slc4a11^−/−^* MCEnC passages 10 and 44, *Slc4a11^+/+^* MCEnC passages 11 and 45 and pHCEnC passage 1 were seeded at 1 × 10^5^ cells/mL in 12-well (MCEnC) and 24-well (pHCEnC) plates and cultured to subconfluence. SLC4A11 was knocked down in confluent pHCEnC with siRNA as described above. ATP was extracted by a boiling water method^56^ and measured using a luciferin–luciferase ATP determination kit (Molecular Probes, Eugene, OR, USA).

### Human Corneal Endothelium from Individuals with CHED

The authors followed the tenets of the Declaration of Helsinki in the treatment of the subjects reported herein. This study was approved by the Institutional Review Board at The University of California, Los Angeles (UCLA IRB #11-000020) and was performed after obtaining informed written consent from the parents of affected individuals with CHED who underwent penetrating keratoplasty.

### Statistical Analysis

All *P* values used in the transcriptomic analysis to identify gene differential expression and pathway enrichment were false discovery rate adjusted *P* values. Statistical analysis, for data other than transcriptomes and patch-clamp recordings, was performed in Prism 7.0 (GraphPad, San Diego, CA, USA) with appropriate statistical tests based on the data structure. Specific statistical tests used for each comparison are indicated in figure legends. Data are presented as mean ± SEM. Statistical significance is denoted as follows in the figures: \**P* < 0.05, \*\**P* < 0.01, \*\*\**P* < 0.001 and \*\*\*\**P* < 0.0001.

## Results

### CHED Corneal Specimens Suggest That *SLC4A11* Mutations Are Not Associated with Decreased Mutant SLC4A11 Expression

Corneal specimens from two individuals with CHED were examined with light microscopy and immunohistochemistry. One of the two individuals (*SLC4A11^Mu/Mu^*) demonstrated compound heterozygous mutations in *SLC4A11* (NM_032034) (c. [473_480delGCTTCGCCinsC; 2623C>T], p.[R158PfsX3; Arg875*]) and the other (*SLC4A11*^Mu/WT^) demonstrated a single heterozygous *SLC4A11* coding region mutation (c.2146C>G, p.Pro716Ala). Both corneal specimens demonstrated a significantly thickened Descemet membrane and an attenuated aberrant CEnC layer with cytoplasmic inclusions present in some cells (Fig. 1A). Immunofluorescence staining for SLC4A11 protein in the corneal endothelium was performed in conjunction with the use of an antibody against an ER marker, protein disulfide isomerase (PDI), to investigate whether these *SLC4A11* mutations lead to protein misfolding and retention in the ER, as previously reported.^5,6,50^ There was no apparent difference in SLC4A11 protein staining intensity between the two specimens from the individuals with CHED, in which the staining localized to the cellular membrane and did not colocalize with PDI, and seven healthy controls (Fig. 1B).

**Figure 1.**
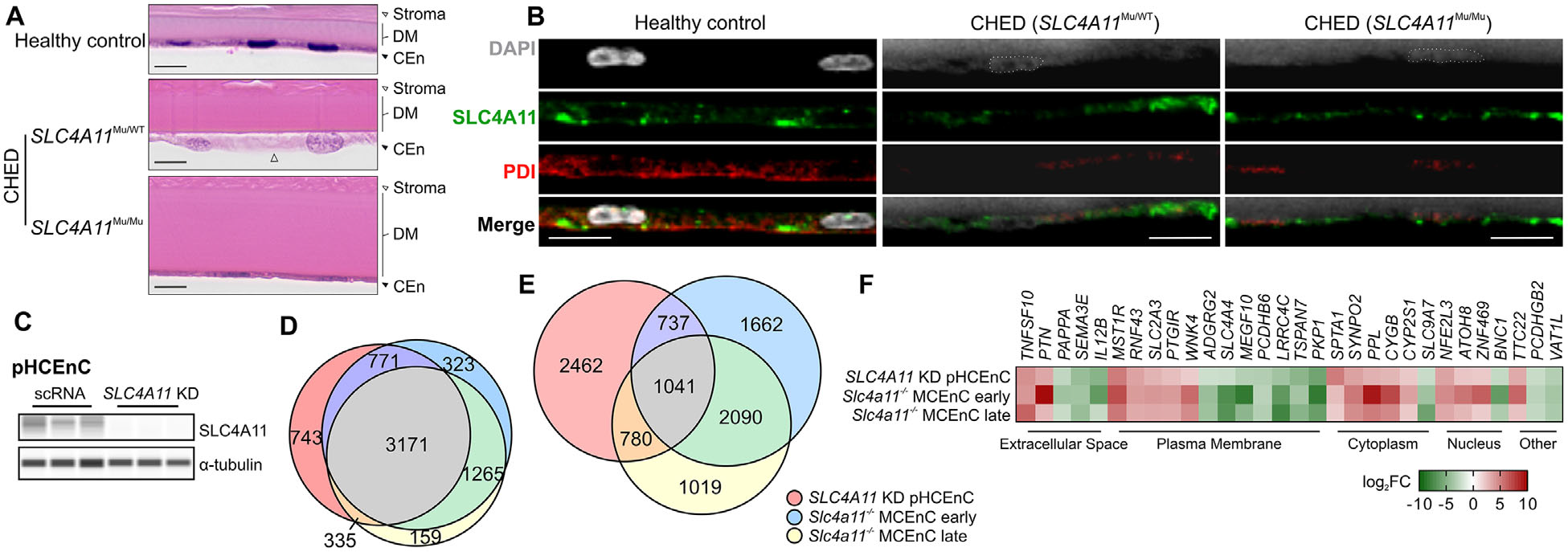
SLC4A11 deficiency leads to transcriptome alteration in corneal endothelium. (**A**) Histopathologic examination of control and CHED corneas demonstrating Descemet membrane (DM) thickening and corneal endothelial cell (CEnC) alteration or attenuation in corneas from individuals with CHED (H&E stain). Δ, cytoplasmic inclusion. *Scale bar*: 10 μm. (**B**) Representative images of immunofluorescence staining for SLC4A11 (*green signal*) and the ER marker PDI (*red signal*) in corneal endothelium of two individuals with CHED and healthy control. Nucleus were stained with 4’,6-diamidino-2-phenylindole (DAPI; *gray signal*). *Scale bar*: 10 μm. (**C**) Western blot analysis for SLC4A11 in pHCEnC treated with siRNA against *SLC4A11* (*SLC4A11* KD) and scRNA for 72 hours. Each lane in the western blot represents an independent primary HCEnC culture established from a unique donor cornea. (**D**) Venn diagram of the endothelial transcriptome in *SLC4A11* KD pHCEnC and *Slc4a11^−/−^* MCEnC early and late passages indicating the number of differentially expressed genes (DEG) in each of the sample sets. (**E**) Venn diagram of the endothelial transcriptome in *SLC4A11* KD pHCEnC and *Slc4a11^−/−^* MCEnC early and late passages indicating the number of DEG in the same direction in each of the sample sets. (**F**) Heat map of top 30 most highly differentially expressed genes shared among *SLC4A11* KD pHCEnC and *Slc4a11^−/−^* MCEnC early and late passages. Genes are clustered based on cellular localization of the gene product.

### SLC4A11/Slc4a11 Reduction Induces Corneal Endothelial Transcriptome Changes

Next, to mimic the loss of SLC4A11 function in CHED, we knocked down SLC4A11 in pHCEnC (Fig. 1C) and utilized an immortalized MCEnC cell line from a *Slc4a11^−/−^* mouse. We then performed transcriptomic analysis of pHCEnC and MCEnC with normal and reduced levels of SLC4A11 and Slc4a11, respectively. Based on the shared corneal endothelial phenotypes between individuals with CHED and the *Slc4a11^−/−^* mouse, we compared the transcriptomes from *SLC4A11* KD pHCEnC and early and late passage MCEnC derived from *Slc4a11^−/−^* mice.^23^ Both early and late MCEnC passages were included in the analysis so that transcriptomic changes due to the loss of Slc4a11 could be differentiated from the transcriptomic changes introduced by prolonged culture. A comparison of the DEG identified from each sample set revealed 3171 genes that were consensually differentially expressed across three sample sets (Fig. 1D), of which 1041 genes were differentially expressed in the same direction (Fig. 1E). Over a third of the 30 most highly DEG (Fig. 1F) have been previously demonstrated to play important functional roles in the cornea and/or have been associated with corneal diseases, including *SEMA3E*,^9^ *MST1R*,^57^ *RNF43*,^58^ *SLC2A3*,^2^ *WNK4*,^59^ *SLC4A4*,^26^ *CYGB*,^60^ *SLC9A7*,^1,61^ *ZNF469*,^62–65^ *BNC1*^66^ and *TTC22*.^67^

### Loss of SLC4A11 Leads to Generalized Inhibition of Cellular Metabolism

Comparison of enriched canonical pathways in *Slc4a11* KD pHCEnC, *Slc4a11^−/−^* MCEnC early (passage 6) and *Slc4a11^−/−^* MCEnC late (passage 39) sample sets identified a shared generalized inhibition (defined by negative activation *z*-score) of multiple metabolic pathways that were interconnected via intermediate metabolites (Figs. 2A, 2B). A generalized decrease in the expression of enzyme-encoding genes in these pathways was observed (Figs. 2C–2I), a finding that was confirmed for selected genes from each pathway using qPCR (Fig. 2J).

**Figure 2.**
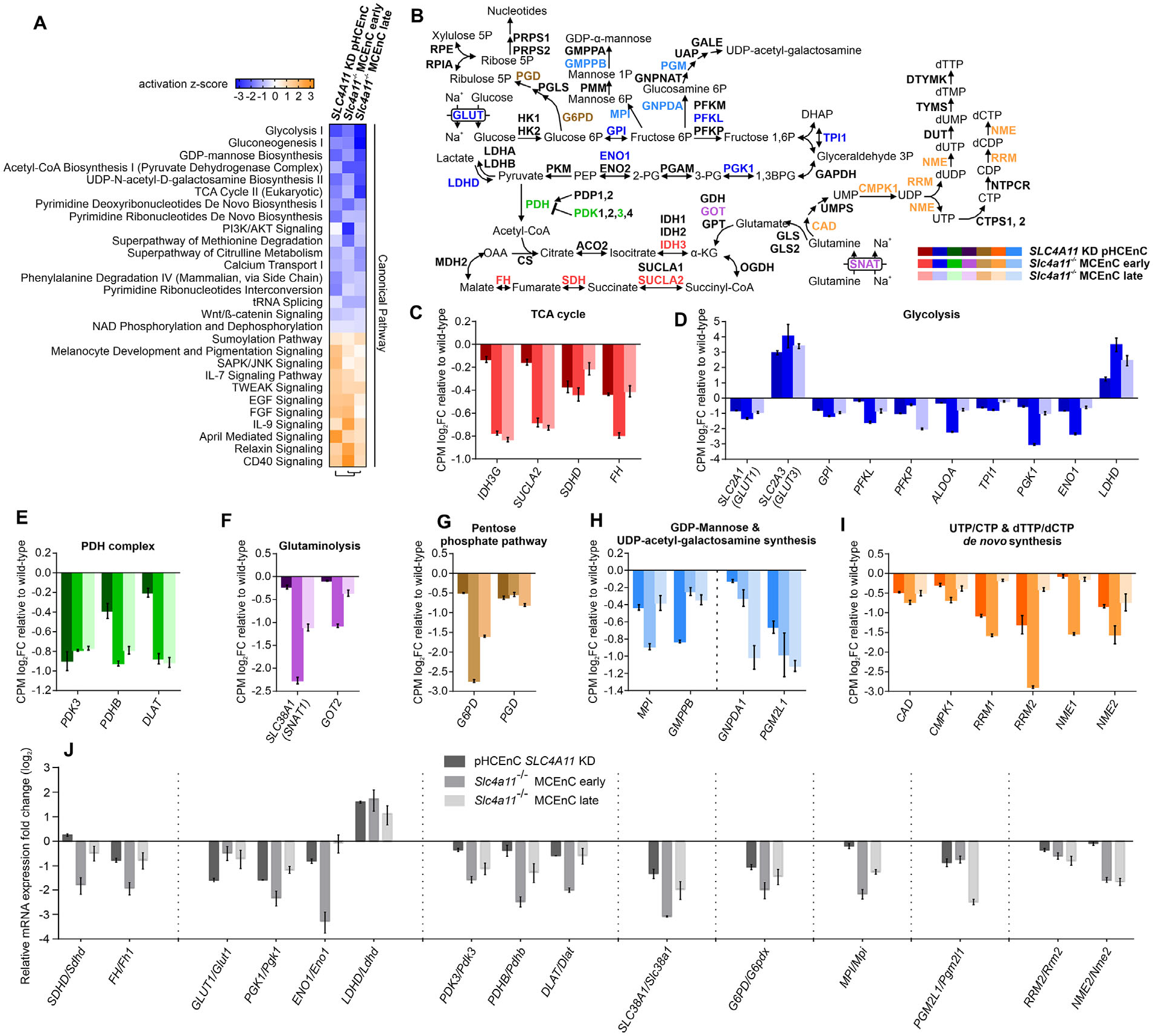
Inhibition of multiple metabolic pathways in SLC4A11-deficient corneal endothelial cells. (**A**) Heat map showing consensually enriched IPA canonical pathways from comparison of transcriptomes of *SLC4A11* KD pHCEnC and *Slc4a11^−/−^* MCEnC early and late passages (sorted by mean activation *z*-score). (**B**) Schematic illustration of the crosstalk among identified inhibited metabolic pathways. Differentially expressed genes encoding key enzymes are color coded using the same colors as used in **C** to **I**. Shown are transcript levels of differentially expressed enzyme-coding genes involved in the following: (**C**) Citric acid cycle (tricarboxylic acid cycle, TCA). *IDH3G,* isocitrate dehydrogenase 3 (NAD^+^) gamma; *SUCLA2,* succinate-CoA ligase ADP-forming beta subunit; *SDHD,* succinate dehydrogenase complex subunit D; *FH*, fumarate hydratas. (**D**) Glycolysis. *GLUT1/3*, glucose transporter type 1 and 3; *GPI*, glucose-6-phosphate isomerase; *PFKL*, phosphofructokinase, liver type; *PFKP*, phosphofructokinase, platelet; *ALDOA*, aldolase, fructose-bisphosphate A; *TPI1*, triosephosphate isomerase 1; *PGK1,* phosphoglycerate kinase 1; *ENO1,* enolase 1; *LDHD,* lactate dehydrogenase. (**E**) Acetyl-CoA biosynthesis/pyruvate dehydrogenase (PDH) complex. *PDK3*, pyruvate dehydrogenase kinase 3; *PDHB*, pyruvate dehydrogenase E1 beta subunit; *DLAT*, dihydrolipoamide *S*-acetyltransferase. (**F**) Glutaminolysis. *SNAT1*, system N amino acid transporter 1; *GOT2*, glutamic-oxaloacetic transaminase 2. (**G**) Pentose phosphate shunt. *G6PD,* glucose-6-phosphate dehydrogenase; *PGD,* phosphogluconate dehydrogenase. (**H**) GDP-mannose synthesis. *MPI,* mannose phosphate isomerase; *GMPPB,* GDP-mannose pyrophosphorylase B in UDP-acetyl-galactosamine synthesis; *GNPDA1,* glucosamine-6-phosphate deaminase 1; *PGM2L1*, phosphoglucomutase 2 like 1. (**I**) Pyrimidine ribonucleotide (UTP/CTP) and pyrimidine deoxyribonucleotide (dTTP/dCTP) de novo synthesis. *CAD*, carbamoyl-phosphate synthetase 2, aspartate transcarbamylase, and dihydroorotase; *CMPK1*, cytidine/uridine monophosphate kinase 1; *RRM1*, ribonucleotide reductase catalytic subunit M1; *RRM2*, ribonucleotide reductase regulatory subunit M2; *NME1,* NME/NM23 nucleoside diphosphate kinase 1; *NME2,* NME/NM23 nucleoside diphosphate kinase 2. (**J**) Selected differentially expressed genes from each pathway listed above were validated by qPCR in separate RNA isolations from *SLC4A11* KD pHCEnC and *Slc4a11^−/−^* MCEnC early and late passages. Genes from different pathways are clustered and separated by dashed lines.

### Altered Expression of Ion Channels and Transporters Impairs Transport Function of Corneal Endothelium

IPA biological function enrichment analysis identified “transport of molecule” as the top inhibited function (negative *z*-score) shared between the transcriptomes of the pHCEnC, MCEnC early and MCEnC late samples sets (Fig. 3A). Because SLC4A11 is an electrogenic NH_3_:H^+^ co-transporter,^15^ we performed single cell recordings of the membrane potential of *Slc4a11^+/+^* and *Slc4a11^−/−^* MCEnC, in which the resting membrane potential is not statistically different (Fig. 3B), in response to extracellularly perfused 10 mM NH_4_Cl. Although exposure to NH_4_Cl induced a +12.7 mV (95% CI, +5 to +20 mV) depolarization in *Slc4a11^+/+^* MCEnC, likely due to the NH_3_:H^+^ permeability provided by Slc4a11, exposure to NH_4_Cl induced a −9.50 mV (95% CI, −19.3 to −0.5 mV) hyperpolarization in *Slc4a11^−/−^* MCEnC (Figs. 3C, 3D).

**Figure 3.**
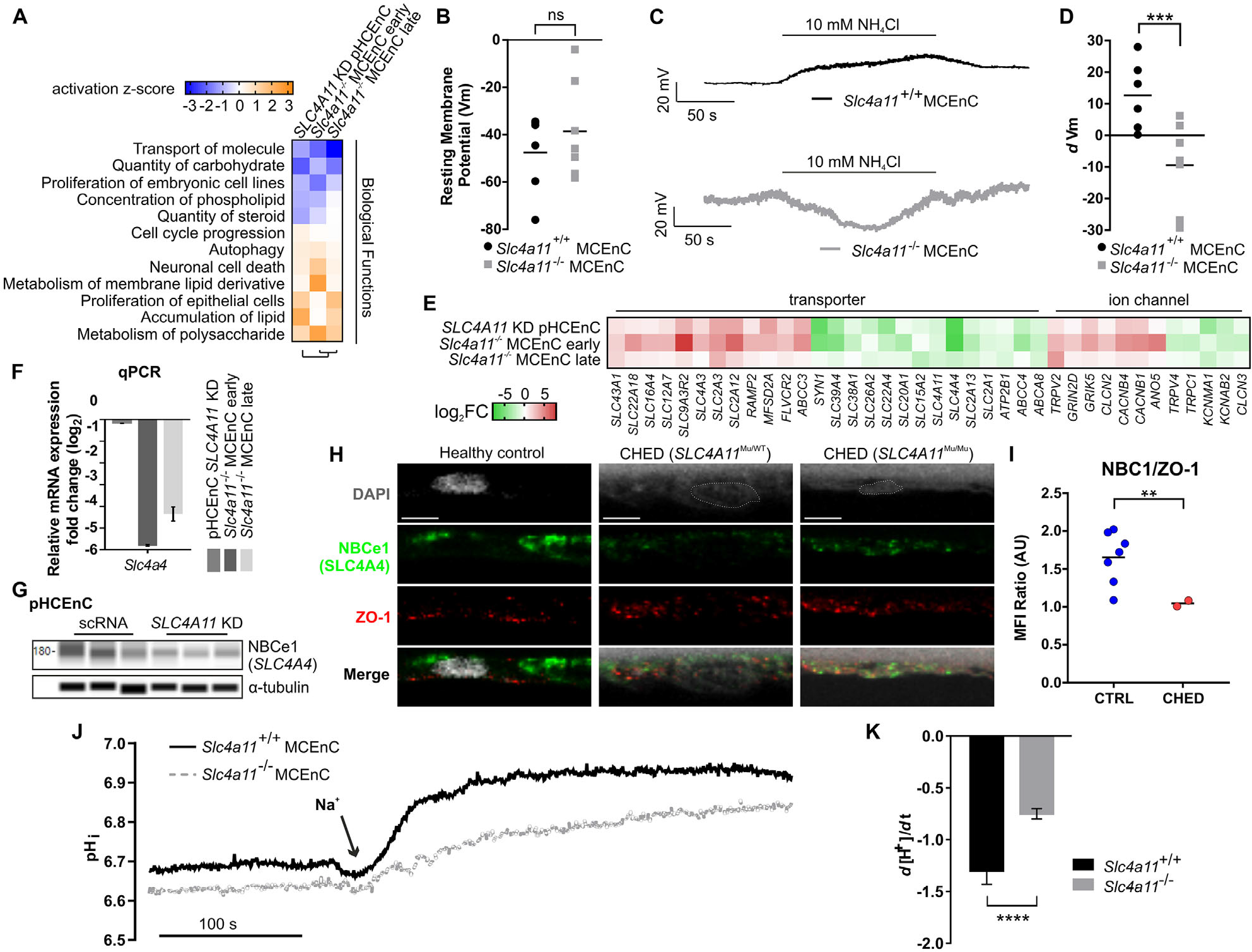
SLC4A11 deficiency impacts corneal endothelial ion and solute transport function. (**A**) Heat map showing consensually enriched IPA biological function pathways from comparison of transcriptomes of *SLC4A11* KD pHCEnC and *Slc4a11^−/−^* MCEnC early and late passages (sorted by mean activation *z*-score). (**B**) Scatterplot of resting membrane potential (Vm) in *Slc4a11^+/+^* (*n* = 6) and *Slc4a11^−/−^* (*n* = 7) MCEnC. Two-tailed paired-samples *t*-test, *P* = 0.408. (**C**) Representative trace of current-clamped single cell recording during 10 mM NH_4_Cl superfusion of *Slc4a11*^+/+^ and *Slc4a11^−/−^* MCEnC. (**D**) Scatterplot of membrane potential changes (dVm) in *Slc4a11*^+/+^ (*n* = 6) and *Slc4a11^−/−^* (*n* = 7) MCEnC in response to NH_4_Cl superfusion. Monte Carlo resampling, two-tailed paired-samples *t*-test, difference between genotype dVm *=* –22.2 (–34.8, –10) mV, *P* = 0.0069. (**E**) Heat map showing common differentially expressed genes encoding ion channel and transporter proteins in *SLC4A11* KD pHCEnC and early and late *Slc4a11^−/−^* MCEnC. (**F**) The differential expression of Na^+^-HCO_3_^−^ transporter (NBCe1, encoded by *SLC4A4*) mRNA was validated by qPCR in separate RNA isolations from *SLC4A11* KD pHCEnC and *Slc4a11^−/−^* MCEnC. (**G**) Western blot for NBCe1 in *SLC4A11* KD pHCEnC and scRNA pHCEnC control showing decreased NBCe1 protein level in *SLC4A11* KD pHCEnC. (**H**) Representative images of immunofluorescence staining for NBCe1 (*green signal*) and tight junction Zonula occludens-1 (ZO-1) (*red signal*) in corneal endothelium of two individuals with CHED and healthy control. Nuclei were stained with DAPI (*gray signal*). *Scale bar*: 5 μm. (**I**) Scatterplot of mean fluorescence intensity (MFI) ratio of NBCe1 over ZO-1 in corneal endothelium of two individuals with CHED and of seven healthy controls. Two-tailed unpaired *t*-test with Welch’s correction, *P* = 0.0031. (**J**) Representative trace of pH_i_, response in *Slc4a11^−/−^* and *Slc4a11^+/+^* MCEnC to the addition of extracellular Na^+^ in HCO_3_^−^-containing solution. (**K**) Bar graph of the rate of intracellular [H^+^] change (*d*[H^+^]/*d*t) in *Slc4a11^+/+^* (*n* = 6) and *Slc4a11^−/−^* (*n* = 8) MCEnC. Two-tailed unpaired *t*-test, *P* < 0.0001.

To investigate the nature of this depolarization to hyperpolarization shift resulting from loss of Slc4a11, we examined the list of 1041 genes differentially expressed in the same direction in the *SLC4A11* KD pHCEnC, *Slc4a11^−/−^* MCEnC early and MCEnC late sample sets to identify ion channels and transporters. We identified 12 genes encoding ion channels and 26 genes encoding transporters were either upregulated or downregulated, including SLC4A11 (Fig. 3E, Table). The electrogenic Na^+^-HCO_3_^−^ cotransporter (NBCe1, *SLC4A4*), which plays an essential role in the CEnC pump function,^26^ was downregulated fourfold in *SLC4A11* KD pHCEnC and >100-fold in *Slc4a11^−/−^* MCEnC (early and late passage). The downregulation of SLC4A4/Slc4a4 was verified by qPCR in *SLC4A11* KD pHCEnC and *Slc4a11^−/−^* MCEnC (Fig. 3F), by western blot in *SLC4A11* KD pHCEnC (Fig. 3G) and by immunofluorescence staining for NBCe1 (SLC4A4) in the corneal endothelium of two individuals with CHED (Fig. 3H), which showed decreased NBCe1 expression compared to controls (Fig. 3I).

Next, functional measurement of Na^+^-dependent HCO_3_^−^ transport in *Slc4a11^−/−^* MCEnC showed reduced Na^+^-HCO_3_^−^ co-transporter activity when compared to *Slc4a11^+/+^* MCEnC (Fig. 3J), consistent with reduced NBCe1 expression. In Figure 3J, pH_i_ was maintained at a low value when *Slc4a11^−/−^* and *Slc4a11^+/+^* MCEnC were perfused with a 28.3 mM bicarbonate (HCO_3_^−^) solution that was Na^+^ free. When switched to a bicarbonate solution containing Na^+^, pH_i_ increased as NBCe1 started to move the weak base HCO_3_^−^ inward using the Na^+^ inward transmembrane electrochemical gradient. We determined the initial slope of this pH_i_ rise to serve as an indirect measure of Na^+^-HCO_3_^−^ cotransport activity and observed a reduced Na^+^-dependent pH, rise in *Slc4a11^−/−^* MCEC compared with *Slc4A11^+,+^* MCEnC (Figs. 3J, 3K).

### Mitochondria Dysfunction Leads to Reduced ATP Production

Dilated mitochondria is a characteristic electron microscopy finding in CHED corneal endothelium, suggestive of mitochondrial involvement in the pathogenesis.^68^ Correspondingly, “mitochondria dysfunction” was the top enriched canonical pathway in the pHCEnC and MCEnC early and late sample sets (ranked by *P* value, Fig. 4A). Additionally, in the list of 1041 genes differentially expressed in the same direction in the pHCEnC and MCEnC early and late sample sets, genes encoding proteins involved in the mitochondria electron transport chain, mediating mitochondrial ATP flux, import machinery and translation machinery showed a generalized decreased expression (Figs. 4B, 4C).^69^ Because transcriptomic analysis also suggested that “autophagy” was activated (Fig. 3A) in *SLC4A11* KD pHCEnC and *Slc4a11^−/−^* MCEnC, we examined the differential expression of *STX17/Stx17,* which encodes SNARE protein Syntaxin 17 (STX17), a mitophagy initiator facilitating the removal of dysfunctional mitochondria.^70,71^ *STX17/Stx17* was upregulated in the *SLC4A11* KD pHCEnC and *Slc4a11^−/−^* MCEnC transcriptomes (Fig. 4D), a finding that was validated by qPCR in *SLC4A11* KD pHCEnC and *Slc4a11^−/−^* MCEnC (Fig. 4E), by western blot in *SLC4A11* KD pHCEnC (Fig. 4F) and by immunofluorescence in CHED corneal endothelium, which demonstrated increased STX17 staining intensity and co-localization with mitochondria marker COX4 compared with healthy controls (Fig. 4G). Significantly decreased COX4 expression was observed in CHED corneal endothelium, indicating reduced mitochondrial density (Figs. 4G, 4I) and resulting in increased STX17 abundance relative to the mitochondrial density (STX17/COX4) (Fig. 4H). Staining CHED endothelium with another mitochondrial marker, cytochrome *c*, also showed consistently decreased staining intensity (Figs. 4J, 4K).

**Table.** Differentially Expressed Genes Encoding Ion Channels and Transporters in *SLC4A11* KD pHCEnC, *Slc4a11^−/−^* MCEnC Early, and MCEnC Late Sample Sets

|  | Upregulated | Downregulated |
| --- | --- | --- |
| Ion channels |  |  |
| Cl<sup>-</sup> channels | *CLCN2, ANO5* | *CLCN3* |
| Ca<sup>2+</sup> channels | *TRPV2, CACNB1, CACNB4* | *TRPV4, TRPC1* |
| K<sup>+</sup> channels | — | *KCNAB2, KCNMA1* |
| Transporters |  |  |
| NH<sub>3</sub>:H<sup>+</sup> transporter | — | *SLC4A11* |
| Na<sup>+</sup>-HCO<sub>3</sub><sup>-</sup> cotransporter | — | *SLC4A4* |
| Ca<sup>2+</sup>-ATPase | — | *ATP2B1* |
| Glucose transporter | *SLC2A3, SLC2A12* | *SLC2A1, SLC2A13, SLC15A2* |
| Glutamine transporter | — | *SLC38A1* |
| Cl<sup>-</sup>/HCO<sub>3</sub><sup>-</sup> exchanger | *SLC4A3* | — |
| ATP-binding cassette transporters | *ABCC3* | *ABCA8, ABCC4* |
| K<sup>+</sup>-Cl<sup>-</sup> cotransporter | *SLC12A7* | — |
| Na<sup>+</sup>/H<sup>+</sup> exchanger regulator | *SLC9A3R2* | — |
| Na<sup>+</sup>-HPO<sub>3</sub><sup>-</sup> cotransporter | — | *SLC20A1* |
| Lactate transporter | *SLC16A4* | — |
| Na<sup>+</sup>-carnitine cotransporter | *SLC22A18* | *SLC22A4* |
| SO<sub>4</sub><sup>2-</sup> transporter | — | *SLC26A2* |

**Figure 4.**
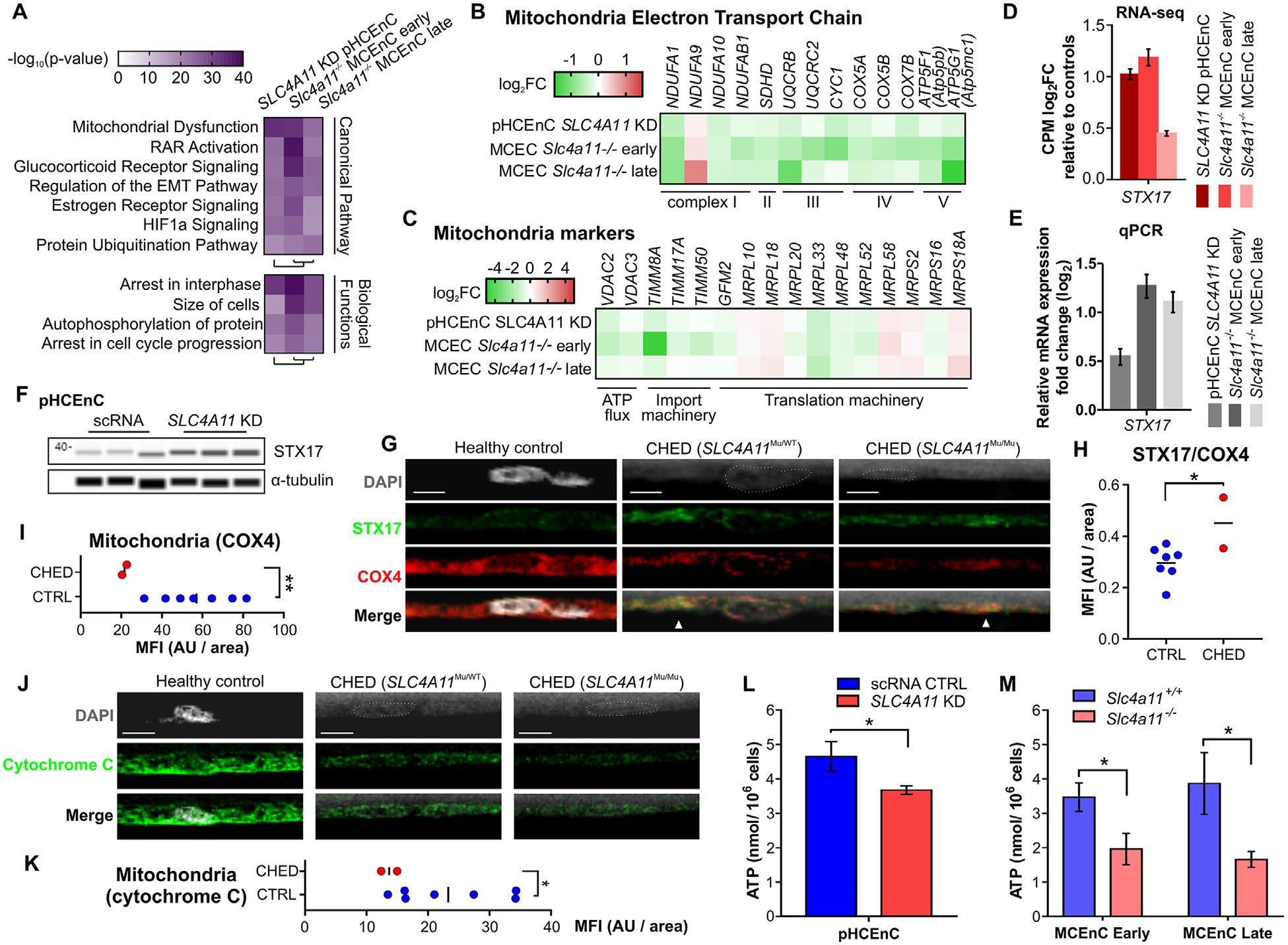
Mitochondria dysfunction in SLC4A11-deficient corneal endothelium. (**A**) Heat map of commonly enriched canonical and biological function pathways in transcriptomes of *SLC4A11* KD pHCEnC and early and late *Slc4a11*^−/−^ MCEnC. (**B**) Transcript levels of differentially expressed enzyme coding genes involved in mitochondria electron transport chain. (**C**) Transcript levels of differentially expressed protein coding genes that serve as mitochondrial functional markers. (**D**) Bar graph of transcriptome data showing transcript-level fold change of *STX17* in *SLC4A11* KD pHCEnC and *Slc4a11*^−/−^ MCEnC compared to controls. (**E**) Bar graph of qPCR showing mRNA expression-level fold change of *STX17* in separate RNA isolations from *SLC4A11* KD pHCEnC and *Slc4a11* MCEnC compared to controls. (**F**) Western blot of STX17 protein level in *SLC4A11* KD pHCEnC and in scRNA pHCEnC controls. (**G**) Representative images of immunofluorescence staining for STX17 (*green signal*) and mitochondria marker COX4 (*red signal*) in corneal endothelium of two individuals with CHED and healthy control. Nuclei were stained with DAPI (*gray signal).* Co-localization of STX17 (*green*) and COX4 (*red*) was seen in CHED corneal endothelium (▲, *yellow*). *Scale bar*: 5 μm. (**H**) Scatterplot of MFI ratio of STX17 over COX4 in corneal endothelium of two individuals with CHED and of seven healthy controls. Two-tailed unpaired *t*-test with Welch’s correction, *P* = 0.0483. (**I**) Scatterplot of MFI of COX4 in corneal endothelium of two individuals with CHED and of seven healthy controls. Two-tailed unpaired *t*-test with Welch’s correction, *P* = 0.0018. (**J**) Representative images of immunofluorescence staining for cytochrome *c* (*green signal*) in corneal endothelium of two individuals with CHED and of seven healthy controls. Nucleus were stained with DAPI (*gray signal*). *Scale bar*: 5 μm. (**K**) Scatterplot of MFI of cytochrome *c* in corneal endothelium of two individuals with CHED and of seven healthy controls. Two-tailed unpaired *t*-test with Welch’s correction, *P* = 0.0304. (**L**) Bar graph summary of intracellar ATP levels measured in *SLC4A11* KD pHCEnC and scRNA pHCEnC controls. One-tailed unpaired *t*-test, *P* = 0.0281 (*n* = 6). (**M**) Bar graph summary of intracellar ATP levels measured in early passage (two-tailed unpaired *t*-test, *P* = 0.034, *n* = 6 each) and late passage (*P* = 0.039, *n* = 6 each) *Slc4a11^+/+^* and *Slc4a11^−/−^* MCEnC.

Given the evidence indicating mitochondrial dysfunction and reduced mitochondrial density in the setting of reduced SLC4A11 expression, we hypothesized that these would result in insufficient ATP energy supply to maintain the Na^+^-K^+^-ATPase-driven corneal endothelial pump and the cornea edema that characterizes CHED. Thus, we performed a direct measurement of the steady-state ATP levels in *SLC4A11* KD pHCEnC and *Slc4a11^−/−^* MCEnC, which revealed reduced ATP concentrations compared to control scRNA pHCEnC and *Slc4a11^+/+^* MCEnC, respectively (Figs. 4L, 4M).

### Transcriptomic Changes Associated with Loss of SLC4A11 Are Mediated by Activation of AMP-Activated Protein Kinase–p53 Pathway

To identify the upstream signaling pathway responsible for the observed transcriptomic changes in *SLC4A11* KD pHCEnC and *Slc4a11^−/−^* MCEnC, we performed upstream regulator prediction in IPA, which identified p53 (encoded by the *TP53* gene) as the top candidate transcription factor (Fig. 5A). Western blot analysis in *SLC4A11* KD pHCEnC demonstrated increased Ser15 phosphorylation of p53 compared to scRNA pHCEnC controls (Fig. 5B), indicative of post-translational activation of p53 transcriptional activity. Similarly, in *Slc4a11^−/−^* MCEnC, we observed increased Ser18 phosphorylation of p53 (corresponding to Ser15 of human p53) in *Slc4a11^−/−^* MCEnC late passage, although this was not observed in *Slc4a11^−/−^* MCEnC early passage (Fig. 5C). However, there was increased total p53 levels in both *Slc4a11^−/−^* MCEnC early and late passages (Figs. 5C, 5D), indicative of transcriptional activation of p53.

**Figure 5.**
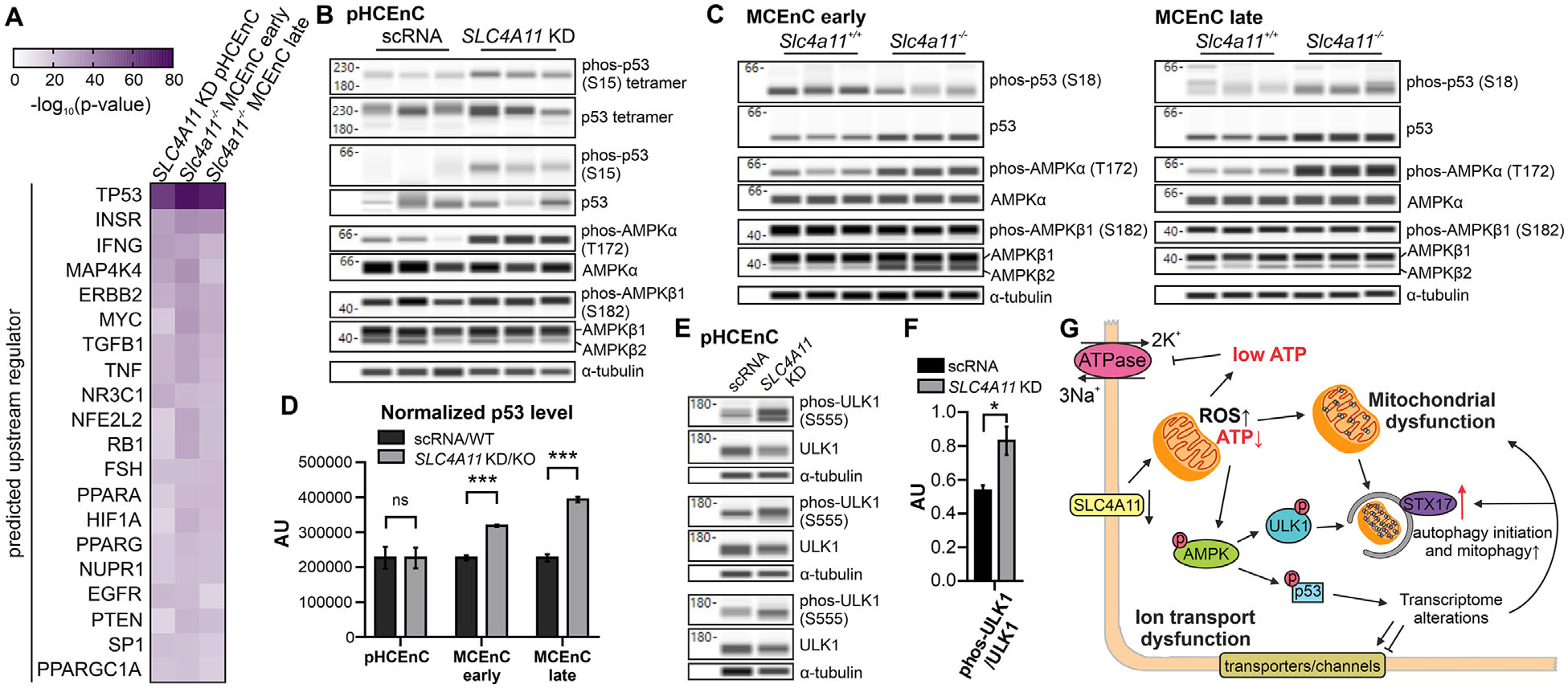
Activation of AMPK-p53 pathway in SLC4A11-deficient corneal endothelium. (**A**) Heat map of shared predicted upstream regulators of transcriptomes of *SLC4A11* KD pHCEnC and early and late *Slc4a11^−/−^* MCEnC (sorted by mean adjusted *P* value). (**B**) Western blot analysis of *SLC4A11* KD pHCEnC and scRNA controls for Ser15 phosphorylated p53 (phos-p53, S15), total p53, Thr172 phosphorylated AMPK*α* (phos-AMPK*α*, T172), total AMPK*α*, Ser182 phosphorylated AMPK*β* (phos-AMPK*β*1, S182), and total AMPK*β*1 and *β*2. (**C**) Western blot analysis of early and late *Slc4a11^+/+^* and *Slc4a11^−/−^* MCEnC for Ser18 phosphorylated p53 (phos-p53, S18), total p53, Thr172 phosphorylated AMPK*α* (phos-AMPK*α*, T172), total AMPK*α*, Ser182 phosphorylated AMPK*β* (phos-AMPK*β*1, S182), and total AMPK*β* 1 and *β*2. (**D**) Bar graph summary of western blot densitometry results for normalized total p53 level in *SLC4A11* KD pHCEnC versus scRNA pHCEnC (two-tailed unpaired-samples *t*-test (*P* > 0.99)), in early passage *Slc4a11^−/−^* versus *Slc4a11^+/+^* MCEnC (*P* = 0.00027) and in late passage *Slc4a11^−/−^* versus *Slc4a11^+/+^* MCEnC (*P* = 0.00021). (**E**) Western blot analysis of *SLC4A11* KD pHCEnC and scRNA control for Ser555 phosphorylated ULK1 (phos-ULK1, S555) and total ULK1. (**F**) Bar graph summary of western blot densitometry results from biological triplicates of *SLC4A11* KD pHCEnC and scRNA pHCEnC controls for the ratio of phosphorylated ULK1/total ULK1 (phos-ULK1/ULK1). Two-tailed unpaired *t*-test, *P* = 0.030. (**G**) Schematic summary of the downstream molecular events as a result of SLC4A11 deficiency in corneal endothelial cells.

We then sought to identify the kinase responsible for the Ser15 (Ser18 in mouse) phosphorylation and transcriptional activation of p53 in *SLC4A11* KD pHEnC and *Slc4a11^−/−^* MCEnC. Given the observed ATP depletion in *SLC4A11* KD pHEnC and *Slc4a11^−/−^* MCEnC, as well as the reported capacity of the cellular ATP sensor AMP-activated protein kinase (AMPK) to mediate Ser15 (Ser18 in mouse) phosphorylation and transcriptional activation of p53, we investigated the potential role of AMPK.^72–74^ In the setting of a decreased ATP-to-adenosine diphosphate (ADP) or ATP-to-AMP ratio, the AMPK catalytic α subunit will be phosphorylated at Thr172, whereas phosphorylation of the regulatory *β*1 subunit at Ser182 is not dependent upon cellular ATP levels.^75^ In *SLC4A11* KD pHCEnC and *Slc4a11^−/−^* MCEnC, we observed increased Thr172 phosphorylation of AMPK*α* and no change in Ser182 phosphorylation of AMPK*β*1 (Figs. 5B, 5C). Examination of another downstream substrate of AMPK, Unc-51 like autophagy activating kinase 1 (ULK1), showed increased phosphorylation (Ser555) in *SLC4A11* KD pHCEnC compared with scRNA pHCEnC (Figs. 5E, 5F).

## Discussion

In this manuscript, we utilized primary human CEnC, a CEnC cell line from *Slc4a11^−/−^* mice^23^ and corneal specimens from individuals with CHED to investigate the causes of CEnC dysfunction that characterize each of the *SLC4A11-* associated corneal endothelial dystrophies (CHED, FECD, and Harboyan syndrome). Although several previous reports have elucidated possible roles for mitochondrial uncoupling, ER unfolded protein response, oxidative stress, and apoptosis in SLC4A11-deficient human and mouse cell lines,^9,20,21,30,50,76^ none utilized primary human CEnC, and only one report examined CHED patient corneal endothelium, in which increased oxidative stress was demonstrated.^20^ We identified ATP depletion in CEnC with reduced SLC4A11, in both transient (72 hours) *SLC4A11* knockdown in pHCEnC and permanent *Slc4a11* knockout in MCEnC. The reduced CEnC ATP levels provide a proposed pathogenesis for the CEnC dysfunction that characterizes the *SLC4A11*-associated corneal endothelial dystrophies. The fact that ATP depletion and ATP-sensor AMPK*α* activation were detected within 72 hours of *SLC4A11* knockdown in pHCEnC suggests that energy shortage from the loss of SLC4A11 is likely among the initial steps in the pathogenesis of *SLC4A11*-associated corneal endothelial dystrophies.

Although the prevailing hypothesis regarding the pathogenesis of CHED is that the majority of *SLC4A11* mutations result in protein misfolding and retention in the ER,^5,6,50,51^ we provide preliminary evidence that mutant SLC4A11 protein is not retained in the ER of the corneal endothelium in CHED. In addition, immunostaining of corneal endothelium from two individuals with CHED did not show a reduced level of the SLC4A11 mutant protein, consistent with a previous report that CHED corneal endothelium does not demonstrate reduced SLC4A11 expression at the mRNA level.^20^ Instead, our data, which we recognize is derived from a very limited number of observations, support the alternative hypothesis that identified *SLC4A11* mutations affect the ion transport function of SLC4A11 protein in the corneal endothelium.^17,52,53^

We used a comparative transcriptomics approach based on the observation of phenotypic similarities between CHED and the *Slc4a11^−/−^* mouse corneal phenotype.^77,78^ This approach is based on the high degree of gene orthology between mouse and human and the organ-dominated hierarchical clustering observed across mammals on real gene expression data.^79^ Such a comparative transcriptomic approach enables the differentiation of transcriptome alterations attributable to the loss of SLC4A11/Slc4a11 in pHCEnC and MCEnC from confounding biological or technical factors associated with each cell preparation, including differences in primary cell isolation and passaging, siRNA treatment, cell line immortalization, and cell culture media.^80^ With this approach, we identified generalized inhibition of multiple metabolic pathways, as well as mitochondria dysfunction in both *SLC4A11* KD pHCEnC and *Slc4a11^−/−^* MCEnC. We also identified the reduced expression of several ion channels and transporters, including NBCe1, and a decrease in Na^+^-dependent HCO_3_^−^ transport activity, which was in contrast to a previous functional examination of HCO_3_^−^ transport in *Slc4a11^−/−^* MCEnC that did not reveal any difference in comparison to *Slc4a11^+/+^* MCEnC.^23^ The discrepancy is likely because previous experiments were performed in a Na^+^-rich, HCO_3_^−^-free solution with subsequent reintroduction of HCO_3_^−^, whereas we performed the experiment in a Na^+^-free, HCO_3_^−^-rich solution with subsequent reintroduction of Na^+^. While a previous report provided an estimate of HCO_3_^−^ transport activities in *Slc4a11^−/−^* MCEnC,^23^ we provided an estimate of Na^+^-dependent HCO_3_^−^ transport activities. We attributed the observed transcriptomic changes in *SLC4A11* KD pHCEnC and *Slc4a11^−/−^* MCEnC to activation of the AMPK–p53 pathway. Although post-translational activation (phosphorylation) of p53 was observed with transient *SLC4A11* knockdown in pHCEnC, transcriptional activation (upregulation) of p53 was observed with permanent *SLC4A11* knockout in MCEnC, both attributed to AMPK*α* activation.^73^ The observation that ATP depletion and AMPK activation occur within 72 hours after SLC4A11 knockdown suggests that they are likely to be among the initial cellular events in response to SLC4A11 deficiency.

In summary, we postulate that SLC4A11 deficiency leads to CEnC dysfunction primarily through decreased generation of ATP via glutaminolysis to fuel the Na^+^/K^+^-ATPase-driven endothelial pump. The decreased ATP levels also result in the activation of AMPK and its downstream substrates, p53 and ULK1, leading to transcriptome alterations and increased mitophagy, respectively. Similarly, given the role of SLC4A11 in preventing oxidative damage, the loss of SLC4A11 leads to increased mitochondrial ROS production,^30^ subsequent mitochondria dysfunction, and increased mitophagy (Fig. 5G). This proposed pathogenesis supports the use of the *Slc4a11^−/−^* mice, in which the alterations resulting from Slc4a11 depletion mirror those observed in *SLC4A11* KD pHCEnC, as a model for the *SLC4A11-* associated corneal endothelial dystrophies and indicates a favorable translational potential for therapeutic approaches shown to be efficacious in the *Slc4a11^−/−^* mice.

## Acknowledgments

Supported by National Eye Institute Grants (1R01EY022082, AJA; P30EY000331 and R01EY029817, APS); the Walton Li Chair in Cornea and Uveitis (AJA); the Stotter Revocable Trust (SEI Cornea Division); an unrestricted grant from Research to Prevent Blindness (AJA); a National Institute of Diabetes and Digestive and Kidney Diseases grant (1R01DK077162, IK); the Allan Smidt Charitable Fund (IK); Ralph Block Family Foundation (IK); and Knights Templar Eye Foundation Career Starter Grant (WZ).

Disclosure: **W. Zhang,** None; **R. Frausto,** None; **D.D. Chung,** None; **C.G. Griffis,** None; **L. Kao,** None; **A. Chen,** None; **R. Azimov,** None; **A.P. Sampath,** None; **I. Kurtz,** None; **A.J. Aldave,** None

