## Supplemental Materials for "Energy Shortage in Human and Mouse Models of *SLC4A11*-Associated Corneal Endothelial Dystrophies"

*Method and Statistical Analysis for Single Cell Patch-clamp Recording*

Single cell recordings were made from isolated single adhered MCEnC cultured on 25-mm glass coverslips to ~10–20% confluence. Coverslips were placed in RC-40LP quick change chambers (Warner Instruments; Hamden CT) and superfused with bicarbonate-buffered Ames' media (equilibrated with 5% $CO_2$/95% $O_2$) heated to 31-33°C with and without 10mM $NH_4Cl$. Membrane voltages were measured using whole-cell patch electrodes in current clamp mode ($I_{hold}$ = 0) as previously described,[1] using an electrode internal solution consisting of (in mM): 125 K-aspartate, 10 KCl, 10 HEPES, 5 NMG-HEDTA, 0.5 $CaCl_2$, 0.5 $MgCl_2$, 1 ATP-Mg, 0.2 GTP-Tris, 2.5 NADPH; pH was adjusted to ~7.3 with NMG-OH and osmolarity was adjusted to ~280 mOsm. MCEnC were superfused with 30–33°C Ames' media buffered with sodium bicarbonate and equilibrated with 5% $CO_2$/ 95% $O_2$, pH~7.4. Ten millimolar of $NH_4Cl$ were added into Ames' media where indicated. Ammonium evoked responses were sampled at 1kHz and digitally low-pass filtered at 50Hz using a 7-pole Butterworth filter. Membrane voltages were averaged over a 10 second window during stable periods of baseline and $NH_4Cl$ application. Ammonium evoked responses were sampled at 1kHz and digitally low-pass filtered at 50Hz using a 7-pole Butterworth filter. Data were reported as mean (95%CI).

Differences between baseline and treatment were calculated using the sample mean for statistical analysis. The normality of the data was assessed visually by quantile-quantile plots and statistically by Shaprio-Wilk tests ($p_{+/+}$ = 0.75; $p_{-/-}$ = 0.32). Due to low sample size (n = 6 and 7 for *Slc4a11*$^{+/+}$ MCEnC and *Slc4a11*$^{-/-}$ MCEnC, respectively), we used

Monte Carlo resampling (bootstrap) and the standard methods to calculate 95% Confidence Intervals and p-values [2]. To determine statistical significance, we considered a false positive rate threshold of $\alpha = 0.05$, and performed a bootstrap paired-sampled t-test using custom scripts in MATLAB (version 2019a, The MathWorks, Natick, 2019). Bootstraps were performed with 10,000 replicates in all cases. Boxplots with dots were used to visualize the measured data. Boxplot whisker methods were calculated using the Tukey 5 number summary method. Briefly, the upper whisker was drawn as Q3 + 1.5 x IQR and the lower whisker was drawn as Q1 - 1.5 x IQR, where IQR is the interquartile range. Datapoints that lie outside of the whisker boundaries are considered outliers. We did not detect any outliers in the data based on this approach.

*Intracellular pH Measurement*

Briefly, pHCEnC and MCEnC were cultured on laminin coated 25-mm diameter glass coverslips. BCECF fluorescence was excited alternately at 500 nm and 440 nm, and the emitted light was collected with a 530 nm filter. Fluorescence ratios (500/440) were obtained at 1 Hz. The cell fluorescence was monitored until a stable $pH_i$ was obtained prior to the start of the experiment. $Na^+$-dependent bicarbonate transport was assayed as follows. The cells were loaded with 10 μM BCECF-AM and bathed in a $Na^+$-free solution containing 140 mM TMACl, 1 mM $CaCl_2$, 1 mM $MgCl_2$, 2.5 mM $K_2HPO_4$, 5 mM dextrose and 5 mM HEPES. The cells were then exposed to a $Na^+$-free bicarbonate solution containing 115 TMACl, 25 $TMAHCO_3$, 1 mM $CaCl_2$, 1 mM $MgCl_2$, 2.5 mM $K_2HPO_4$ and 5 mM dextrose bubbled with $CO_2$ at a pH of 7.4. Following a steady state period, the cells were bathed in a $Na^+$-containing solution with 115 NaCl, 25 $NaHCO_3$, 1 mM $CaCl_2$, 1 mM $MgCl_2$, 2.5 mM $K_2HPO_4$ and 5 mM dextrose bubbled with $CO_2$ at a pH of 7.4 and the change in $pH_i$ was monitored.

**Supplemental Table 1.** Primer sequences used for qPCR

| | species | Forward Sequence (5' -> 3') | Reverse Sequence (5' -> 3') | Amplicon Size |
|---|---|---|---|---|
| *Eno1* | mouse | TGCGTCCACTGGCATCTAC | CAGAGCAGGCGCAATAGTTTTA | 118 |
| *Ldhd* | mouse | CTGAAGGCAGTTGTAGGGAGC | GGAACACCTTGATTGTAGCACAG | 170 |
| *Glut1* | mouse | CAGTTCGGCTATAACACTGGTG | GCCCCCGACAGAGAAGATG | 156 |
| *Pgk1* | mouse | ATGTCGCTTTCCAACAAGCTG | GCTCCATTGTCCAAGCAGAAT | 164 |
| *fh1* | mouse | GAATGGCAAGCCAAAATTCCTT | CGTTCTGTAGCACCTCCAATCTT | 133 |
| *g6pdx* | mouse | CACAGTGGACGACATCCGAAA | AGCTACATAGGAATTACGGGCAA | 103 |
| *Sdhd* | mouse | TGGTCAGACCCGCTTATGTG | GGTCCAGTGGAGAGATGCAG | 128 |
| *Actb* | mouse | GGCTGTATTCCCCTCCATCG | CCAGTTGGTAACAATGCCATGT | 154 |
| *Pdk3* | mouse | TCCTGGACTTCGGAAGGGATA | GAAGGGCGGTTCAACAAGTTA | 133 |
| *Pdhb* | mouse | CGGTGCAGTTGACAGTTCGT | TCTTCCCCAAGCAGAAAAACTTT | 88 |
| *Dlat* | mouse | TCCCTCCGCATCAGAAGGTT | CCAACTGGAACATCTCTGGTC | 217 |
| *Ppia* | mouse | GAGCTGTTTGCAGACAAAGTTC | CCCTGGCACATGAATCCTGG | 125 |
| *Stx17* | mouse | CCAGATCCACAACTGTTGATGG | GCAGCATTTTGGTCTTGAGGAA | 112 |
| *Slc4a4* | mouse | GATGCCACCGACAACATGC | TCAAGATGGTAAGCGGTTGAC | 103 |
| *Mpi* | mouse | CCGCGAGTGTTCCCACTTT | TGTCCCCATCCACAGCTCT | 150 |
| *Pgm2l1* | mouse | ATGACCTGAACTCTAACCTGCT | CCCATTCCGCAAGAGATTTTCAA | 137 |
| *Rrm2* | mouse | TGGCTGACAAGGAGAACACG | AGGCGCTTTACTTTCCAGCTC | 110 |
| *Nme2* | mouse | AGCAGCATTACATCGACCTGA | CATCACTCGGCCCGTTTTCA | 125 |
| *Slc38a1* | mouse | CCTTCACAAGTACCAGAGCAC | GGCCAGCTCAAATAACGATGAT | 127 |
| *FH* | human | GGAGGTGTGACAGAACGCAT | CATCTGCTGCCTTCATTATTGC | 133 |
| *LDHD* | human | TGACTGGTCACCCTGCCA | AGCTCTCCCTTTGCCTTCTG | 132 |
| *GLUT1* | human | ATTGGCTCCGGTATCGTCAAC | GCTCAGATAGGACATCCAGGGTA | 174 |
| *PGK1* | human | GAACAAGGTTAAAGCCGAGCC | GTGGCAGATTGACTCCTACCA | 137 |
| *SDHD* | human | TTGCTCTGCGATGGACTATTCC | CAAGGCATCCCCATGAACAT | 100 |
| *PPIA* | human | TCCTGGCATCTTGTCCAT | TGCTGGTCTTGCCATTCCT | 179 |
| *ACTB* | human | GAAGATCAAGATCATTGCTCCT | TACTCCTGCTTGCTGATCCA | 111 |
| *ENO1* | human | TGGTGTCTATCGAAGATCCCTT | CCTTGGCGATCCTCTTTGG | 126 |

| | | | | |
|---|---|---|---|---|
| *G6PD* | human | CGAGGCCGTCACCAAGAAC | GTAGTGGTCGATGCGGTAGA | 166 |
| *PDK3* | human | CGCTCTCCATCAAACAATTCCT | CCACTGAAGGGCGGTTAAGTA | 156 |
| *PDHB* | human | AAGAGGCGCTTTCACTGGAC | ACTAACCTTGTATGCCCCATCA | 153 |
| *DLAT* | human | CCGCCGCTATTACAGTCTTCC | CTCTGCAATTAGGTCACCTTCAT | 136 |
| *STX17* | human | TCCTTTGACCAGATCCATGACT | CTTGAGGAATTTCAGGTAAGGCA | 107 |
| *SLC4A4* | human | TCTCCAGTGCAAGTAGGATGT | GGTCCTTCTCCGGTTTATCAGA | 114 |
| *MPI* | human | CAGAGGACAAGCCTTATGCAG | GGTGTTTCAACTGAGAGCACTT | 196 |
| *PGM2L1* | human | TGGCTCCGCTGGGATAAGAA | GCAACAAAGACGATCTCGCAG | 96 |
| *RRM2* | human | CACGGAGCCGAAAACTAAAGC | TCTGCCTTCTTATACATCTGCCA | 129 |
| *NME2* | human | CGCACTTTAGTGCCAGGACC | CGGAATCCCTTCTGCTCGAA | 121 |
| *SLC38A1* | human | CACCACAGGGAAGTTCGTAATC | CATCCACGTACCAGGCTGAAA | 152 |

**Supplemental Table 2.** Antibodies Used in Immunofluorescence and Western Blot

| **Primary antibody** | **Host** | **Concentration used** | **Vendor** |
|---|---|---|---|
| anti-SLC4A11 | rabbit | IHC 1:100, Wes 1:25 | Custom-made |
| anti-PDI | mouse | IHC 1:100 | Enzo Life Sciences, SPA-891 |
| anti-α-tubulin | mouse | Wes 1:50 | Cell Signaling, 3873 |
| anti-NBCe1 | rabbit | IHC 1:100, Wes 1:50 | Custom-made |
| anti-STX17 | rabbit | IHC 1:100, Wes 1:50 | GeneTex, GTX120212 |
| anti-COX4 (F-8) | mouse | IHC 1:100 | Santa Cruz, sc-376731 |
| anti-cytochrome C (136F3) | rabbit | IHC 1:100 | Cell Signaling, 4280 |
| Alexa Fluor® 488 F(ab')2 anti-Rabbit IgG | Goat | IHC 1:200 | Invitrogen |
| Alexa Fluor® 568 F(ab')2 Anti-Mouse IgG | Goat | IHC 1:200 | Invitrogen |
| anti-p53 | mouse | Wes 1:50 | Cell Signaling, 2524 |
| anti-phospho-p53 (Ser15) | rabbit | Wes 1:50 | Cell Signaling, 9284 |
| anti-AMPKα | rabbit | Wes 1:50 | Cell Signaling, 5831 |
| anti-phospho-AMPKα (Thr172) | rabbit | Wes 1:50 | Cell Signaling, 2535 |
| anti-AMPKβ | rabbit | Wes 1:50 | Cell Signaling, 4178 |
| anti-phospho-AMPKβ (Ser182) | rabbit | Wes 1:50 | Cell Signaling, 4186 |
| anti- phospho-ULK1 (Ser555) (D1H4) | rabbit | Wes 1:50 | Cell Signaling, 5869 |
| anti-ULK1 (D8H5) | rabbit | Wes 1:50 | Cell Signaling, 8054 |
